# Substantia nigra degradation results in widespread changes in medial zona incerta afferent and efferent connectomics

**DOI:** 10.1101/2021.09.01.458438

**Authors:** Linda H. Kim, Taylor Chomiak, Michelle A. Tran, Stephanie Tam, Claire McPherson, Shane E. A. Eaton, Young Ou, Zelma H. T. Kiss, Patrick J. Whelan

## Abstract

Parkinson’s disease (PD) is a complex disease affecting many facets of movement, especially gait abnormalities such as shuffling and freezing of gait. The nigrostriatal pathways of the basal ganglia are traditionally targeted by existing therapies; however, other pathways may be more relevant to gait, such as the pedunculopontine nucleus and the zona incerta (ZI). The A13 nucleus may be such a target as it has emerged as an area of interest in dopamine motor function. Yet, this area remains understudied compared to other dopamine nuclei, especially in animal models of PD. In 6-OHDA mice, we found a reduction in locomotion in the open field and gait dysfunction during treadmill tests. Medial ZI dopamine cells, containing the A13 nucleus, were preserved following 6-OHDA, in contrast to a marked reduction in substantia nigra pars compacta (SNc) neurons. There was extensive remodelling of the A13 afferent and efferent connectome following nigrostriatal lesions. Afferent input patterns displayed a marked reduction in cross-correlation across brain regions in 6-OHDA mice, while efferent projections showed an increase. In a human PD patient with advanced gait dysfunction we found that the A13 nucleus was preserved, suggesting that remodelling could also occur in humans. This work points to the A13 region as a potential therapeutic target in PD.

**Significance Statement:** Recently it was found that the medial zona incerta projects to the cuneiform nucleus suggesting a parallel dopaminergic projection onto motor regions. Here we investigated the connectome of the A13 region and examined the afferent and efferent projections in normal mice and mice with a unilateral Parkinsonian mouse model. We found that the connectome was reconfigured following nigrostriatal degeneration. This work provides a comprehensive insight into the plasticity in a dopaminergic-rich area of the zona incerta in PD injury models.

## Introduction

Parkinson’s disease (PD) is a complex condition affecting many facets of function, with the most obvious being motor dysfunction. With progressive degeneration of the nigrostriatal dopaminergic pathways, some patients may progress from bradykinesia to postural instability, akinesia, and freezing of gait (1–4). Current treatments such as pharmacological medications, deep brain stimulation (DBS) and physical therapies lead to only partial gait improvements and these remain major problems in advanced stages of PD (5, 6). While subthalamic nucleus (STN) and globus pallidus (GPi) are common DBS targets for PD, alternative targets such as pedunculopontine nucleus (PPN) and the zona incerta (ZI) have been proposed with mixed results in improving postural and/or gait dysfunctions (6–15). Part of the issue with targeting the ZI with DBS strategies is the relative lack of knowledge regarding its downstream anatomical and functional connectivity with motor centres. Data regarding connectivity and its capacity to change during PD is of critical importance. Recently we have discovered that the dopaminergic A13 nucleus located within ZI projects to two areas of the mesencephalic locomotor region (MLR), the PPN and cuneiform nucleus (CUN) (16). This suggests a role for the A13 nucleus in promoting locomotor activity through a dopamine pathway that is parallel to the nigrostriatal pathway. The A13 nucleus is a dopaminergic region located within the medial portion of the rostral and a small medial portion of the dorsal and ventral ZI (17, 18). This region provides dopaminergic input onto dorsolateral periaqueductal grey (dlPAG), superior colliculus (SC), paraventricular nucleus of the thalamus (PVT), central nucleus of the amygdala (CeA), and thalamic reuniens (RE) and has been implicated in a variety of approach and avoidance behaviors (19–29). Of interest is that PD patients often experience freezing of gait in narrow spaces, in the dark, and when they are under time constraints (30). Since the A13 connectome includes projections to motor and visual regions, an interesting possibility is that changes in A13 connectivity following nigrostriatal degeneration could contribute towards compensatory or aberrant connectivity associated with freezing of gait.

While work is not extensive, neurons in the ZI-hypothalamic region, including A13 region, are preserved in advanced PD brains in humans. Recent work in primates and mice (31–34) also appears to show that the A13 is preserved in PD models. However, there is also evidence of plasticity within this region following nigrostriatal denervation. For example, ZI neurons in rats following nigrostriatal denervation have been shown to increase their average firing rates (35). However, it remains unknown if there are any changes in afferent and efferent projections from this region. In PD, there are many pathological- and compensatory-driven changes throughout the neuraxis. Thus, to gain more insight into using the A13 region as a potential alternative target to treat gait dysfunctions in PD, we identified areas of preservation and plasticity within the A13 connectome using whole-brain imaging techniques. We found evidence of a global remodelling of afferent and efferent projections of the A13 region following lesions of the nigrostriatal pathway using a 6-hydroxydopamine neurotoxin (6-OHDA). Furthermore, we demonstrate that the A13 region is preserved in a human patient with advanced gait dysfunction. Overall, these data show that the A13 connectome exhibits distinct patterns of preservation and plasticity across the neuraxis and highlight potential therapeutic pathways in advanced PD. A portion of these data have been published in abstract form (36).

## Results

### Dopaminergic Cells in ZI Are Preserved in PD in human and mouse models

We observed preservation of TH^+^ cells in the A13 region (Fig 1A) within a human PD brain even after a disease duration of 20 years. The clinical diagnosis of PD was confirmed at autopsy by severe depletion of neurons in the SNc bilaterally (Fig 1B) and the LC (Fig 1C), along with the presence of Lewy bodies in the SNc, LC, dorsal motor nuclei of the vagus, brainstem, and cerebral cortex. Clinical features of PD in the patient included freezing of gait, bradykinesia, and rigidity with a Unified Parkinson Disease Rating Scale motor (UPDRS-III) score (37) of 32 off and 11 on medication prior to bilateral subthalamic deep brain stimulator implantation in June 2008. He had good results with sustained benefit such that 5 years post-DBS his best on medication/on stimulation UPDRS-III score was 18. Despite this, his speech worsened over time, as did his gait with multiple falls. He had a sudden death, likely pulmonary or cardiac, at the age of 80, 10 years after DBS surgery and 20 years after PD diagnosis.

**Figure 1.**
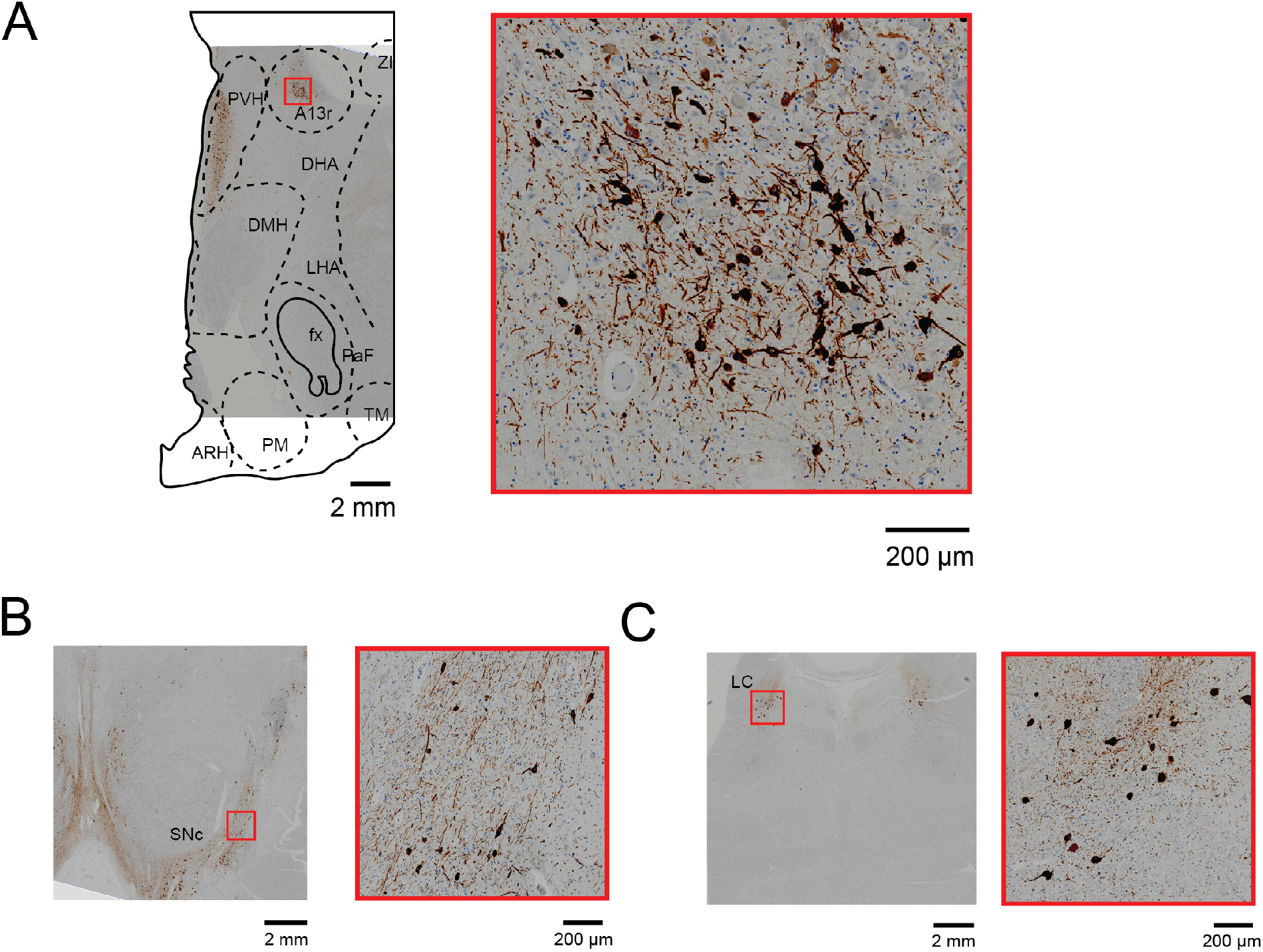
TH^+^ A13 cells are preserved in a patient with Parkinson’s disease. Histological coronal sections indicate TH^+^ immunopositive cells. (A). A13 region is preserved. (B,C). There was an extensive loss of TH^+^ cells in the SNc (B) and the LC (C). *A13r*: A13 rostral, *ZI:* zona incerta, *PVH:* paraventricular nucleus of the hypothalamus, *DMH:* dorsomedial hypothalamus, *LHA:* lateral hypothalamic area, *DHA:* dorsal hypothalamic area, *fx:* fornix, *PaF: parafornical nucleus, ARH:* arcuate nucleus of the hypothalamus, *PM:* posteromedial nucleus, *TM: tuberomamillary nucleus*.

To examine changes in the ZI in greater detail we used a well-validated unilateral 6-OHDA mediated Parkinsonian mouse model (38) (Experimental design - Fig 2). We calculated the percentage of TH^+^ cell loss within the lesioned side normalized to the unlesioned side. We found a significant interaction between the treatment group and brain regions (F(2,12)=5.576, p=0.019, Fig 3A-C). 6-OHDA treated mice showed a significantly greater percentage of TH^+^ cell loss in SNc (−71.98%) compared to VTA (−28.53%) and ZI (−25.70%) (*post-hoc* t(5)=6.075, p=0.005 and t(3)=5.942, p=0.029, respectively). In contrast, sham control mice showed no significant difference in TH^+^ cell density across SNc, VTA and ZI. Thus, similar to that observed in the human Parkinson’s brain, there is remarkable preservation of dopaminergic cells in ZI after nigrostriatal degeneration in the 6-OHDA mouse model of PD.

**Figure 2.**
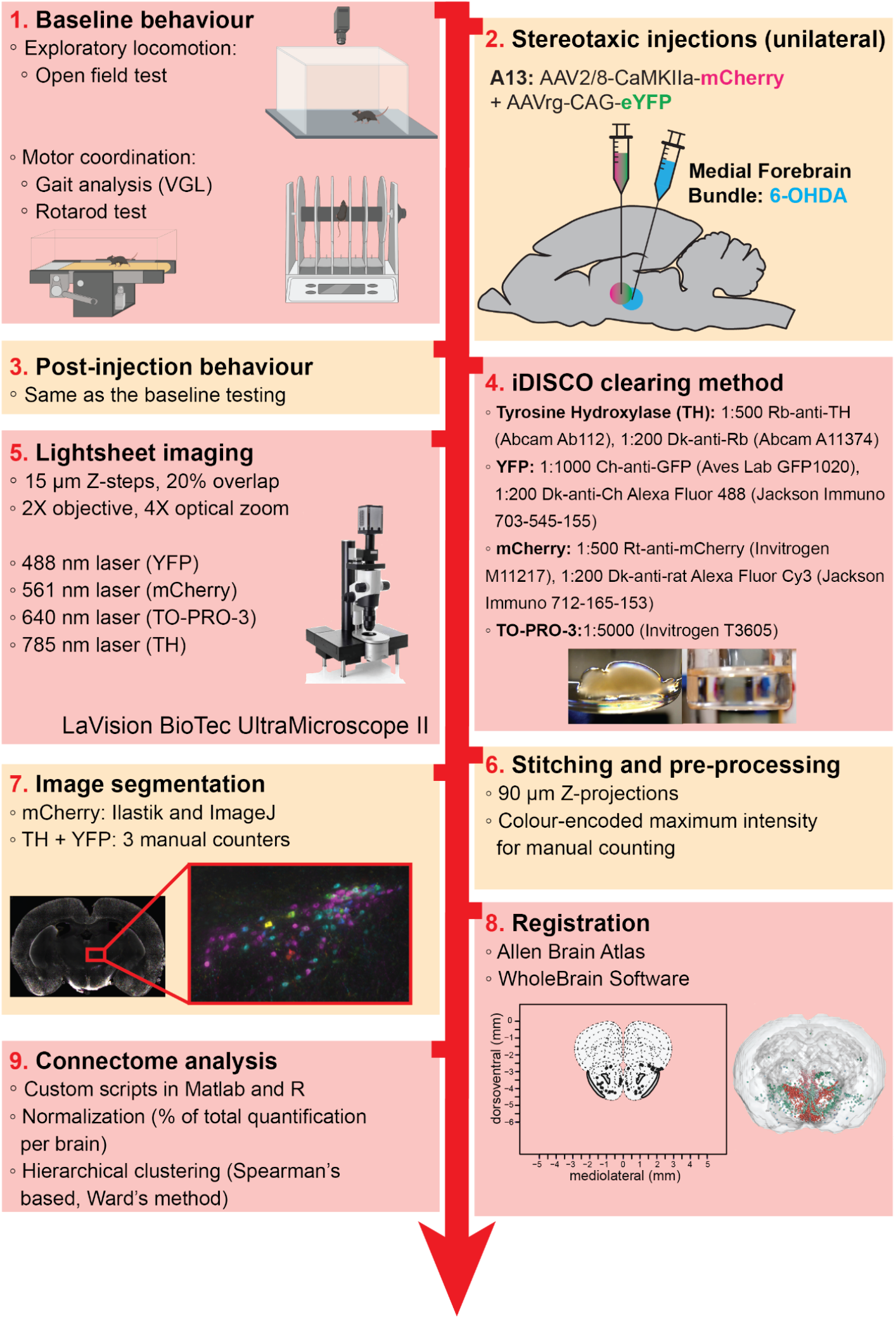
Overview of the methodology used and timeline. Baseline behavior was tested, then a unilateral injection of 6-OHDA or vehicle into the medial forebrain bundle was performed. Anterograde and retrograde injections of viral tracers were directed towards the A13/mZI incerta. Once post-injection behavior was assessed, the brains were harvested and cleared using iDISCO protocols. Lightsheet microscopy was utilized to capture optical coronal slices. Following manual counting, and image segmentation of fibres, the slices were registered using the Allen Brain Atlas and WholeBrain software. Cross-correlation and hierarchical clustering for connectome analysis were performed using R functions.

**Figure 3.**
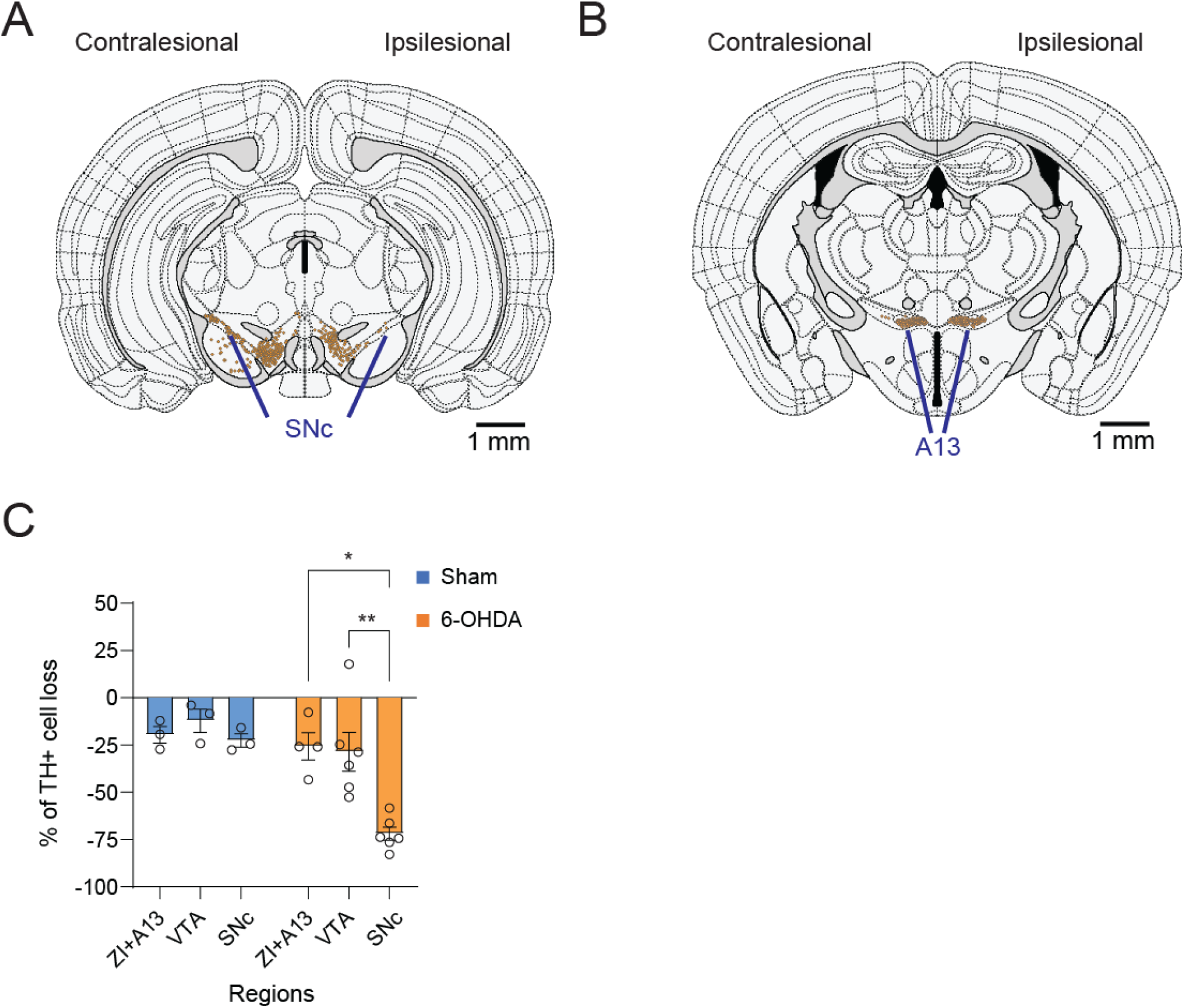
Preservation of TH^+^ A13 cells in Parkinsonian mouse models. Representative slices (450 µm maximum intensity projection) of SNc (AP: -3.08 mm, A) and A13 region (AP: -1.355 mm, B) registered with WholeBrain software. Full 3D brain is available (see Movies S1). There was a lack of TH^+^ SNc cells following 6-OHDA injections at the Medial Forebrain Bundle (MFB) (A). Meanwhile, TH+ VTA cells were preserved bilaterally. In addition, TH^+^ A13 cells were present ipsilesional to 6-OHDA injections (B). When calculating the percentage of TH^+^ cell loss normalized to the unlesioned side, there was a significant interaction between the treatment group and brain regions (C). The 6-OHDA injected group showed a significantly greater percentage of TH^+^ cell loss in SNc than VTA and ZI. In contrast, the sham group showed no significant difference in TH^+^ cell density across SNc, VTA and ZI. * p ≤ 0.05, and * * p ≤ 0.01.

### 6-OHDA mediated Loss of Dopaminergic Cells in SNc Leads to Loss of Exploratory Behaviors and Gait and Sensorimotor Dysfunctions

To examine changes in exploratory behaviour following 6-OHDA mediated nigrostriatal degeneration, we compared locomotor behaviour before and after stereotaxic injections. The change in locomotor behaviour was calculated by the percentage of change observed 3 weeks after 6-OHDA injections normalized to the measurements taken pre-injection for each mouse. 6-OHDA-treated mice displayed a significant loss of exploratory locomotor behaviour in distance travelled, number of locomotor bouts, duration of bouts, and velocity compared to sham mice (Fig 4B-E). 6-OHDA treated mice showed a dramatic decrease in the percentage of locomotor bouts (Mdn=-71.82%, IQR=37.98%) compared to sham (Mdn=3.289%, IQR=45.82%, U=0, p=0.012, one-tailed, Fig 4B). In addition to decreased number of bouts, the duration of bouts was also shortened in the 6-OHDA group (M=-60.79%, SEM=8.04%) versus the sham group (M=-15.06%, SEM=14.21%, t(7)=3.06, p=0.009, one-tailed, Fig 4C). The 6-OHDA group showed a greater velocity loss (Mdn=-71.82%, IQR=3.29%) post-injection compared to sham (Mdn=3.29%, IQR=4.20%, U=0, p=0.012, one-tailed, Fig 4D). Overall, the 6-OHDA group showed a greater loss in total distance travelled post-injection (M=-63.35%, SEM=8.57%) compared to the sham (M=2.60%, SEM=20.05, t(7)=3.632, P=0.004, one-tailed, Fig 4E).

**Figure 4.**
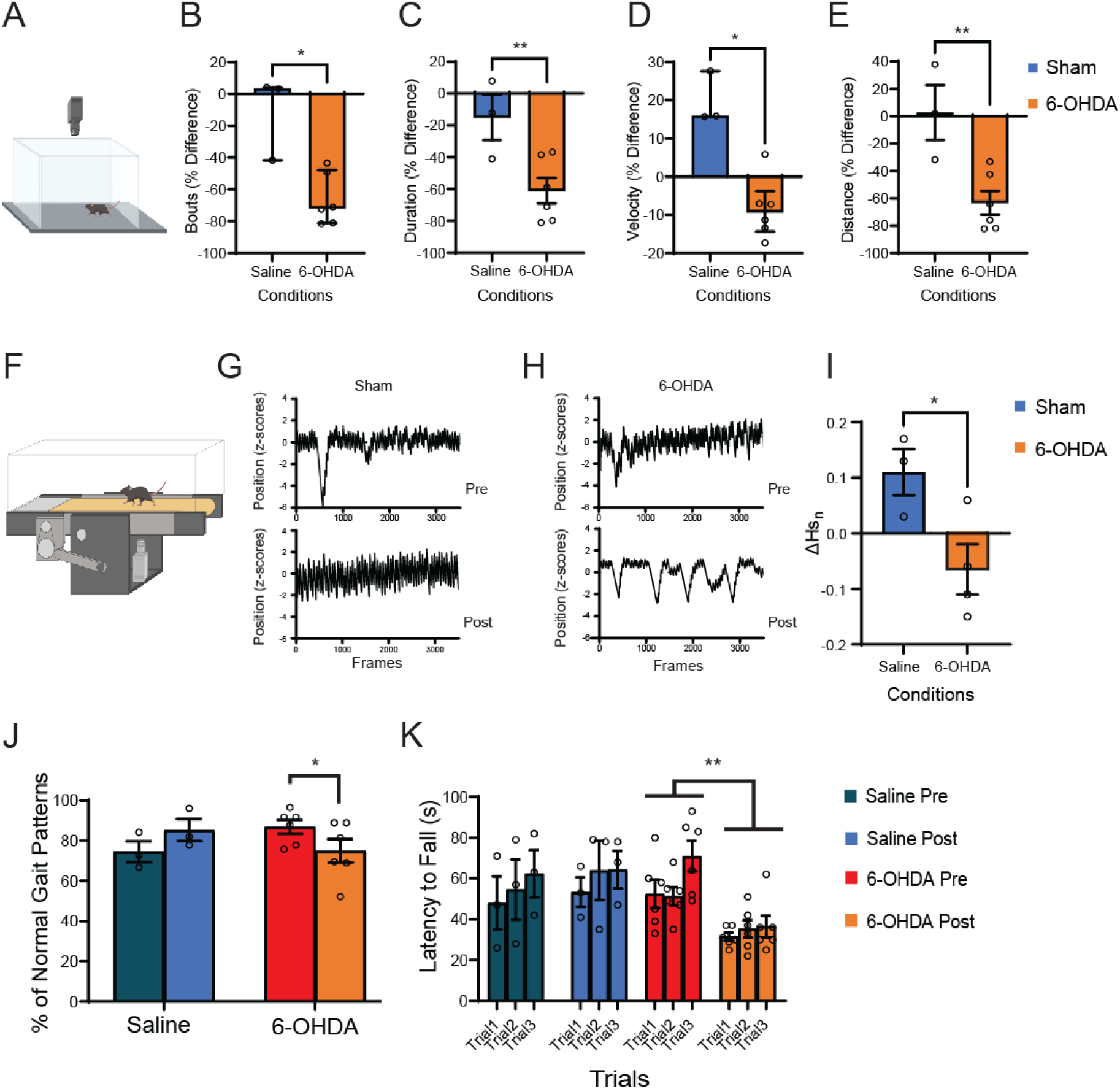
6-OHDA mediated nigrostriatal degeneration led to decreased exploratory behaviour and abnormalities in gait and interlimb coordination. 6-OHDA injected group had significantly greater loss of open field locomotor behaviour (A) from their baseline compared to the sham, in terms of number of locomotor bouts (B), duration of movement (C), total distance travelled (D), and average velocity (E). Next, to determine how well mice could maintain sustained locomotion on a motorized treadmill (F), the contralesional hindpaw positions were tracked using the VGL software (G, H). The 6-OHDA injected group displayed more intermittent locomotor bouts during the post-injection period compared to the shams. The inability of 6-OHDA mice to remain stationary relative to the camera underneath a motorized treadmill was represented by the change in normalized spectral entropy (ΔHs_n_) (39, 40). Higher Hs_n_ is associated with fewer deviations in stationarity. There was a significant decrease in Hsn following a 6-OHDA injection (I). Sham mice did not show any significant change in Hsn. Even when mice could sustain a consistent locomotor bout, 6-OHDA injected mice had a significant reduction in the gait regularity index (% use of normal gait patterns) compared to the sham (J). Rotarod test also revealed a significant decrease in motor coordination following 6-OHDA injections. 6-OHDA group showed significantly shorter latency to fall post-injection than following vehicle injections and pre-injection periods (K). * p ≤ 0.05, and * * p ≤ 0.01.

The reduction in locomotor bouts, distance travelled, and velocity decrease were further investigated to determine the pattern of gait dysfunction, using a motorized treadmill (Fig 4F). Normalized spectral entropy (Hs_n_) was used to assess the inability of 6-OHDA mice to remain stationary relative to the camera underneath the treadmill (39, 40). Hs_n_ provides information about the overall complexity of signal fluctuations; higher values being associated with fewer deviations in stationarity during continuous treadmill running. 6-OHDA mice displayed discontinuous bouts of locomotion more frequently than sham mice (Fig 4G, H), and there was a significant decrease in Hs_n_ following a 6-OHDA injection (M=-0.065%,SEM=0.046%) compared to vehicle injection (M=0.11%, SEM=0.042%, t(5)=2.727, p=0.021, one-tailed, Fig 4I).

Increased variability within sustained locomotor behaviour can be due to many factors, including abnormalities in gait itself. Even when 6-OHDA mice could sustain a consistent locomotor bout where they remain stationary relative to the treadmill camera, their gait became uncoordinated following the 6-OHDA injection (Fig S1). There was a significant difference in the swing to stance duration across all four limbs post-injection in the 6-OHDA group, but not the sham group (time and condition interaction F(1,28)=16.53, p=0.0004, Fig S1A). Gait symmetry between forelimbs to hindlimbs duration was also significantly decreased following 6-OHDA injection in the 6-OHDA group but not within the sham group (time and condition interaction F(1,14)=8.754, p=0.010, *post-hoc* t(14)=4.26, p=0.002, Fig S1B). In contrast to sham mice, 6-OHDA-injected mice displayed significant changes in gait patterning. They showed a significant decrease in use of normal gait patterns (F(1,7)=11.46, p=0.012, Fig 4J). Ab step sequence remained the dominant step sequence pre- and post-injection periods in sham mice. While 6-OHDA mice had normal Ab step sequence dominance pre-injection, Ab dominance was lost following 6-OHDA injection (Step type, condition and time interaction F(5,42)=11.04, p<0.001, Fig S1C). Other gait features such as lengths and durations of stance, swing or stride did not significantly change over time for either group (Fig S1). Finally, during the rotarod test, motor coordination decreased following 6-OHDA injections but not vehicle injections (time and condition interaction F(1,7)=10.863, p=0.013). Bonferroni-adjusted comparisons indicated that 6-OHDA group showed significantly shorter latency to fall (M=34.44 s, SEM=1.51 s) post-injection compared to pre-injection period (M=58.28 s, SEM=6.37 s, p=0.002, Fig 4K). In contrast, latency to fall did not significantly change following vehicle injections in the sham group (p=0.470).

These data indicate that unilateral lesion of SNc DA neurons leads to a robust decrease in exploratory behaviour and subtle abnormalities in gait and interlimb coordination. The next step was to establish the connectome changes that occurred in 6-OHDA compared to sham mice.

### Large-Scale Changes in the A13 connectome Following 6-OHDA Mediated Unilateral Nigrostriatal Degeneration

To examine whether unilateral nigrostriatal degeneration resulted in changes in the organization of the A13 afferent and efferent proportions across the neuraxis, we assessed interregional correlations following 6-OHDA injection compared to vehicle injection. To minimize the influence of experimental variation on the total labelling of neurons and fibres, the afferent cell counts or efferent fibre areas in each brain region were divided by the total number found in a brain to obtain the proportion of total inputs and outputs. For cross-correlation analyses, the data were normalized to a log_10_ value to reduce variability and bring brain regions with high and low proportions of cells and fibres to a similar scale (41). The data across animals were consistent with an average correlation of 0.91 ± 0.02 (Spearman’s correlation, Figure S2).

We first visualized interregional correlations of afferent and efferent proportions for each condition (A13 afferent and efferent proportions of shams, and A13 afferent and efferent proportions of 6-OHDA). Correlation matrices were organized respecting the hierarchical anatomical groups from the Allen Brain Atlas (see Fig S2, *SI Appendix*, and *Materials and Methods* for more details). We observed clear differences in projection patterns between the sham and the 6-OHDA group (Fig 5A-D). Overall, afferents onto the A13 region in sham displayed a higher level of cross-correlation between brain regions compared to 6-OHDA injected mice. In 6-OHDA injected mice, there were 2 distinct clusters of the anti-correlated fraction of inputs from isocortical (motor, sensory, visual, and prefrontal), striatal, and pallidal subregions. These data suggest that afferents from several regions showed a coordinated reduction in afferent density onto the A13 region. But this reduction was not observed for A13 efferents. In marked contrast, the projection patterns of A13 efferents exhibited a higher level of cross-correlation between brain regions following a unilateral nigrostriatal degeneration compared to sham. These data indicate that A13 efferents are in a position to exert greater effects across the neuraxis following nigrostriatal degeneration.

**Figure 5.**
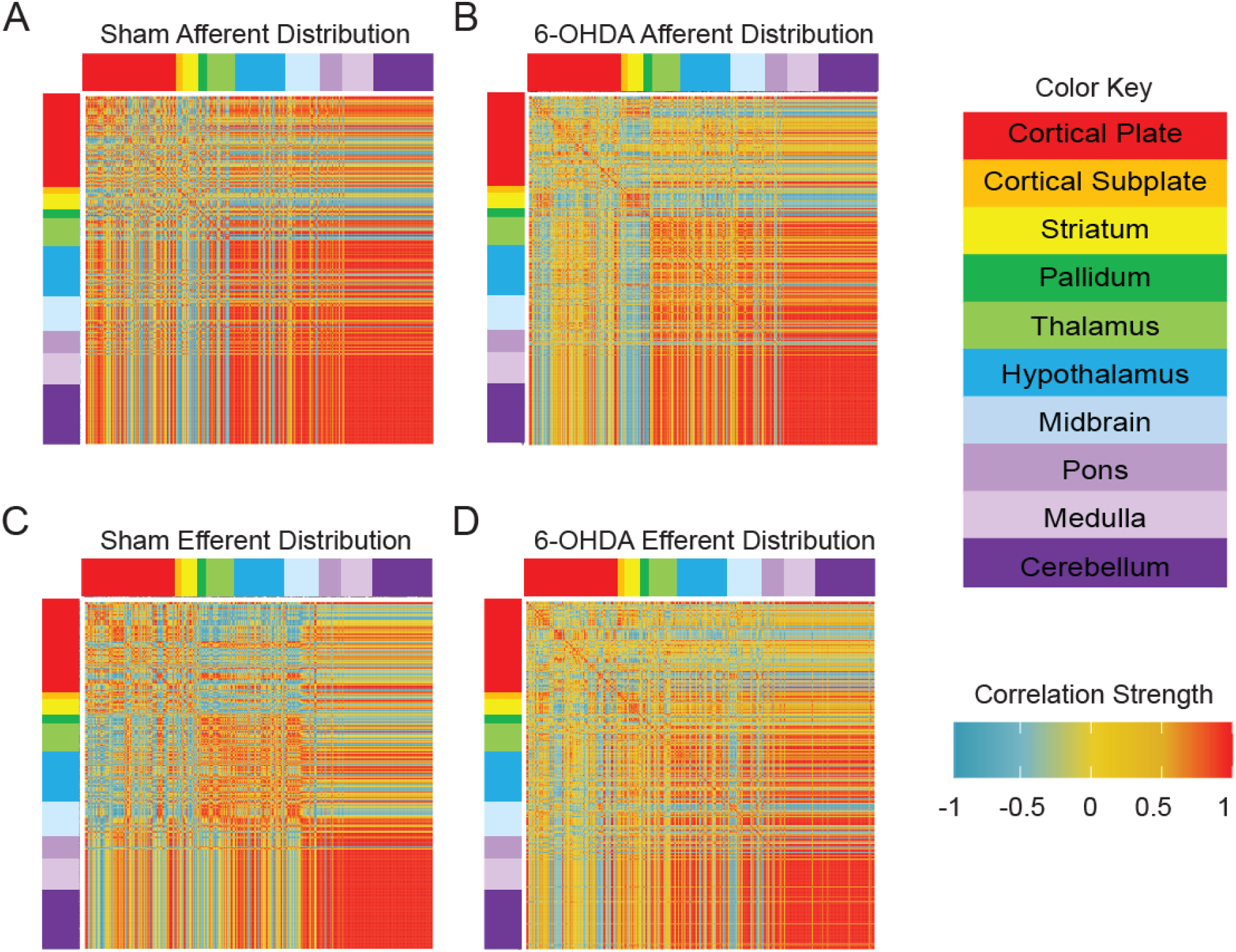
Unilateral nigrostriatal degeneration leads to large-scale changes in the organization of the A13 afferent and efferent distributions across the neuraxis. Interregional correlations of afferent and efferent distributions across the neuraxis identified dissimilar (anti-correlated) regions (see Fig S2, *SI Appendix*, and *Materials and Methods* for more details). The afferent distribution pattern in the sham displayed a higher level of cross-correlation between brain regions (A) than 6-OHDA injected mice (B). Indeed, two distinct bands of anti-correlated afferent regions were identified in the 6-OHDA injected mice. These two bands spanned across the isocortical (motor, sensory, visual, and prefrontal), striatal, and pallidal subregions. In contrast, the projection patterns of A13 efferents displayed a higher level of cross-correlation between brain regions following a unilateral nigrostriatal degeneration (D) compared to sham (C).

### Nigrostriatal Degeneration Leads to Increased Anatomical Modularity

Functional network reorganization and changes in brain modularity of PD patients has been reported previously (42, 43). We hypothesized that similar changes in modular organization of the A13 connectome occurred in our PD mouse models. Using hierarchical clustering of complete Euclidean distance matrices to identify modular organization, we found a modest increase in modularity following nigrostriatal lesioning. The increase in the number of modules that was caused by 6-OHDA induced lesions was independent of the clustering thresholds that were used (Fig 6A,D). The projection patterns of both afferents and efferents of A13 were organized into 3 modules in sham and 4 modules in 6-OHDA treated mice (Fig 6B,C,E,F). In both A13 afferent and efferent networks of 6-OHDA injected mice, hierarchical clustering identified two modules with opposing distribution patterns and two modules with moderate correlations. These increases in modular structuring of the brain are indicative of major structural reorganization of both afferents and efferents of the A13 region.

**Figure 6.**
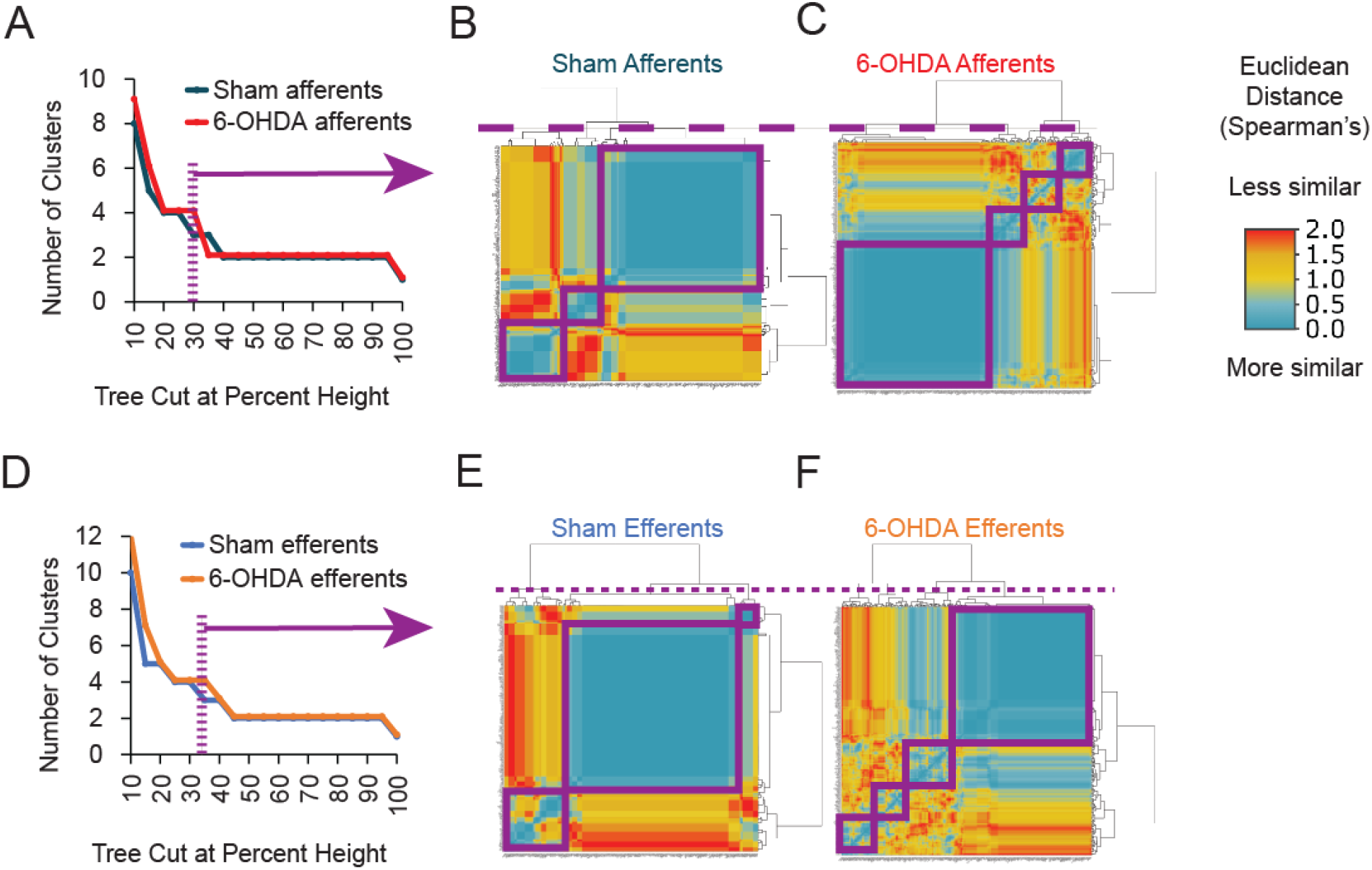
Changes in modular organization of the A13 connectome following a unilateral nigrostriatal degeneration. The hierarchical clustering of complete Euclidean distance matrices based on the distribution of afferents and efferents of A13 across the neuraxis was used to identify the modular organization. In the 6-OHDA injected mice, we found a modest increase in modularity. This increase in the number of modules was independent of the clustering thresholds that were used (A, D). The projection patterns of both afferents and efferents of A13 were reorganized from 3 modules in sham (B, E) to 4 modules in 6-OHDA treated mice (C, F). Increased modularity suggests major structural reorganization of both afferents and efferents of the A13 region.

### Differential Remodelling of A13 Connectome Following 6-OHDA Mediated Nigrostriatal Degeneration

The distributions of A13 connectome in sham served as a basis for an in-depth comparison of preservation and plasticity of A13 afferents and efferents in 6-OHDA mouse models (Fig 7A,E). We observed a global remodelling of A13 afferents and efferents following unilateral nigrostriatal degeneration that was differentially expressed across the neuraxis (Fig 7B, F). As expected, the ipsilesional side showed more areas of downregulation (ipsilesional proportion of total input densities - sham: 63.91% and 6-OHDA mice: 53.81%, overall loss of -10.10% compared to sham, Fig 7D). These downregulated areas were focused within the isocortex and cortical subplate regions (loss of 8.58% and 3.49% of total input density, respectively). 6-OHDA injections also downregulated A13 afferent densities from the striatum, pallidum, thalamus and medulla (loss of 0.81%, 0.27%, 0.71% and 0.08% of total input density, respectively). While 6-OHDA mainly downregulated ipsilesional A13 afferent densities, the hypothalamus (including zona incerta), midbrain, pons, and cerebellum demonstrated increased A13 afferent densities (0.99%, 2.20%, 0.65% and 0.01% increase in proportions of the total input, respectively).

**Figure 7.**
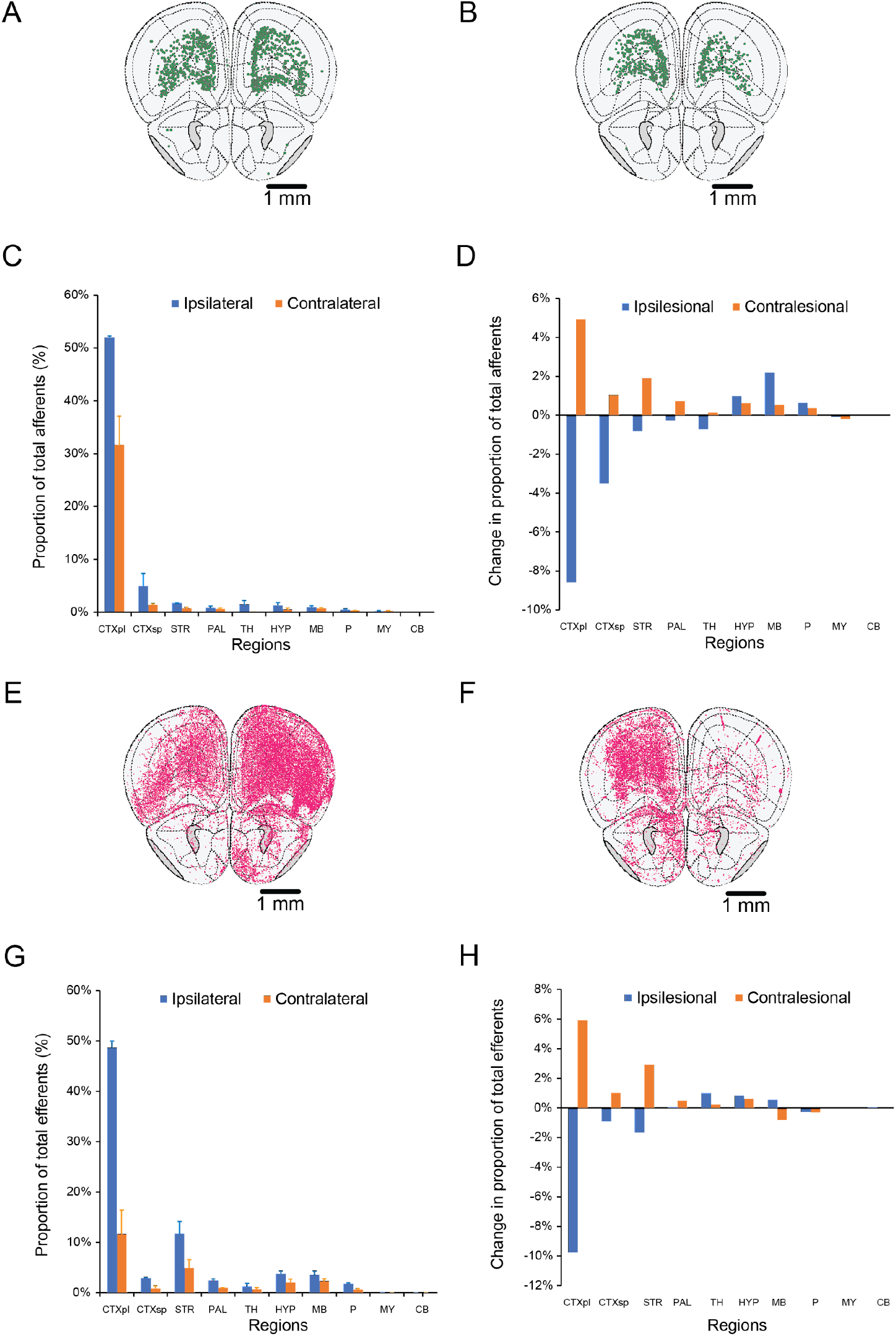
Differential remodelling of A13 connectome following a unilateral nigrostriatal degeneration. The distributions of the A13 connectome in sham mice served as a basis for an in-depth comparison against 6-OHDA mouse models. A. Example slices at rostral areas show changes in intact afferents compared to B. 6-OHDA lesioned animals. Ipsilesional is on the right in figure B. C. Graph showing major brain regions contributing afferents to A13/mZI in intact mice. D. Graph illustrates the change in the proportion of afferents in 6-OHDA compared to intact mice. E. Representative slice showing intact proportions of efferents in intact compared to F. 6-OHDA mice. G. Graph showing major brain regions showing efferents from A13/mZI in intact mice. H. Graph illustrates the change in the proportion of efferents in 6-OHDA compared to intact mice.

The unlesioned, contralesional side showed more upregulated regions across the neuraxis, suggestive of compensatory upregulation in the unilateral 6-OHDA model (contralesional proportion of total input densities - sham: 36.09% and 6-OHDA mice: 46.19%, overall gain of 10.10% in 6-OHDA mice, Fig 7D). The isocortex, striatum, cortical subplate were the top 3 upregulated contralesional regions (gain of 4.93%, 1.92%, and 1.04% of total input density, respectively). Moreover, unlike their counterparts on the lesioned side, which were downregulated, contralesional A13 afferent densities from the pallidum and thalamus were spared and upregulated (0.72% and 0.14% of total input density, respectively). Thus, compensatory upregulation of A13 afferent density from these regions appeared to be lateralized from the unlesioned, contralesional side. Furthermore, bilateral compensatory upregulation of A13 afferents was observed from the hypothalamus (including zona incerta), midbrain and pons. Interestingly, the medulla was the only broad region to show bilateral downregulation of A13 afferent density, with the unlesioned side being affected by 6-OHDA (loss of total input density 0.08% ipsilesional, 0.20% contralesional).

The A13 efferents demonstrated a similar remodelling pattern as above. There were more downregulated areas on the ipsilesional side (ipsilesional proportion of total input densities - sham: 76.02% and 6-OHDA mice: 65.95%, overall loss of -10.07% compared to sham, Fig 7H). However, the downregulation was focused mostly within the isocortical, striatal and cortical subplate regions (loss of 9.74%, 1.67%, and 0.90% of total input density, respectively). Remodelling on the contralesional efferent projection patterns closely followed the changes seen with the afferents, except for projections onto the thalamic and midbrain regions. The A13 efferents onto thalamic regions were bilaterally upregulated (1.00% on the lesioned side and 0.23% on the unlesioned side). In addition, the A13-midbrain efferents were upregulated ipsilesionally (0.56%) and downregulated contralesionally (−0.80%).

## Discussion

PD is now well recognized to affect multiple brain structures beyond the nigrostriatal projection. Here we focused on the medial portion of rostral ZI which contains dopaminergic and GABAergic neurons involved in the multimodal control of locomotor activity. Our work demonstrates clear changes in locomotor behaviour following a unilateral 6-OHDA induced lesion of the substantia nigra. Over four weeks, there was a remarkable change in both the afferent and efferent A13 region connectome, despite the preservation of TH^+^ ZI cells. The afferent projections were particularly downregulated ipsilesionally, while contralesionally projecting afferents showed upregulation. In contrast, efferent projections showed less downregulation onto the cortical subplate regions and bilateral upregulation of thalamic and hypothalamic efferents.

### Impact of nigrostriatal lesion afferent source on A13 connectome

The large-scale changes of A13 connectome that we observed in our study align with previous studies that explored the interconnection between ZI and basal ganglia and the effects of lesioning afferent sources. Previous tracing studies in rats have shown that dorsal and ventral ZI projections, predominantly glutamatergic, preferentially target the SN and PPT of the basal ganglia (44). Both regions show significant degeneration in Parkinson’s disease and animal models (45–48). Lesioning of afferent sources could lead to transneuronal plasticity and contribute further to the symptoms of Parkinson’s disease. ZI cells are hyperactive in Parkinsonian models (35). Parvalbumin expression, dominant within the ZI and mostly colocalizing with GABA (17, 49, 50), is significantly reduced within the mediolateral extent of ZI following a unilateral 6-OHDA mediated nigrostriatal degeneration (51). Interestingly, Nissl staining revealed little to no cell death, suggesting that the cells changed expression patterns. Recently, several limbic regions across the amygdala and hypothalamus have shown significant upregulation of TH signal intensity in MPTP-treated Parkinsonian mice (34). In our work, the interconnection between the dopaminergic A13 region and hypothalamus (including zona incerta) are upregulated bilaterally in the 6-OHDA injected mice. Meanwhile, limbic cortical subplate afferents onto the A13 region were downregulated ipsilesionally to a greater extent than A13 efferents projecting to cortical subplate regions. Downregulation of limbic A13 afferents may contribute to deficits in emotional processing in Parkinson’s disease (52). It remains unclear whether the global change of the A13 connectome is directly related to loss of afferent sources or secondary to abnormal activity from afferents induced by 6-OHDA.

### Possible role of preserved A13 in PD

Previous work in ZI has shown chemoarchitectural diversity among the four ZI sectors and differences in interconnections with the basal ganglia network (see reviews: (12, 17)). Rostral ZI can be differentiated from others through dopaminergic neurons (A13 region) and interconnections with the cerebral cortex, especially with the cingulate cortex (53). The sham group in our work showed similar strong interconnections with the cerebral cortex. These cortical-A13 afferents and efferents were also downregulated following nigrostriatal degeneration. Moreover, following the nigrostriatal degeneration, afferent projection patterns from the isocortical (motor, sensory, visual, and prefrontal) areas remain anti-correlated with the rest of the neuraxis. In contrast, the efferent projection patterns show greater cross-correlation across the neuraxis. This pattern of reorganization suggests that the A13 region can maintain the biased isocortical input while its output becomes more distributed. Our findings suggest that executive control of behavioural response, which descending ZI efferents could modulate, remain intact following nigrostriatal degeneration. ZI has been proposed to be an integrative node for functionally diverse somatosensory and viscerosensory inputs (54) with widespread interconnections among the four sectors (55). A13-ZI interconnections are upregulated on the contralesional side and A13-hypothalamic interconnections upregulated bilaterally in 6-OHDA injected mice.

PD is a heterogeneous disease with motor and nonmotor symptoms directly resulting from neuronal loss or aberrant connectivity of surviving neurons. The nigrostriatal degeneration is believed to produce the core motor symptoms of PD (bradykinesia, rigidity, and tremor) and contribute to cognitive, mood, and sleep dysfunctions. It is thought that the core motor symptoms appear when a critical threshold of loss in striatal terminals (>70%) and SNc cells (>50%) has been reached (56–60). However, preclinical symptoms are masked by early compensatory mechanisms, and clinical deterioration is likely due to inadequate compensatory mechanisms or degeneration of non-dopaminergic neurons. Our experimental PD model represents an advanced PD state with ⪆60% SNc degeneration where changes in the A13 connectome could have been involved in early and active compensation but not necessarily be able to overcome motor deficits. Although compensatory mechanisms are thought to be local, involving sensitization of remaining spared striatal terminals, increased dopamine synthesis and release, increased availability of extracellular dopamine (61), and increased sprouting of striatal dopaminergic fibres (62–64), non-striatal dopaminergic systems, such as the nigropallidal pathway, have also been implicated (65). The limbic basal ganglia network is relatively preserved and could provide a large source of collaterals to adjacent denervated sensorimotor and associative territories (64). Likewise, preserved A13 efferents targeting the basal ganglia network may provide compensatory dopaminergic innervation with collateralization and increasing availability of extracellular dopamine.

While core motor symptoms respond well to dopamine replacement therapy early on, the disease often progresses and leads to gait and balance impairments that only partially respond. Freezing of gait (FOG) is a gait dysfunction characterized by sudden and brief episodes of being unable to produce forward stepping and a loss of gait automaticity. With the regions involved in stimulus-response habitual control being more susceptible to degeneration (eg. ventrolateral SNc and caudolateral sensorimotor putamen), there is an increased reliance on goal-directed control of movement through the frontocortical-associative basal ganglia network (66, 67). Dopaminergic or DBS therapy may improve freezing of gait, but treatment response is incomplete. Corticoincertal networks that remain intact in PD could ease the burden of increased cognitive demand. Some freezers can overcome freezing of gait by using exteroceptive cues such as visual, auditory and attentional cues to compensate for the deficit of internal cueing and/or proprioceptive integration (68–70). Previous work suggests that the rostral ZI coordinates viscerosensory input with appropriate visceral, arousal, attention, and motor responses. Recent studies using cell type and/or projection-specific approaches to target the rostral ZI region suggest a modulatory role in goal-directed behaviour such as defensive and fear responses (28, 71, 72) and appetitive drive (73, 74). The A13 region provides major dopaminergic innervation onto SC (75) and dorsolateral PAG (16, 24), which have been implicated in modulating orienting and defensive motor responses, respectively (76). Photoactivation of glutamatergic dorsolateral PAG neurons increased occurrences of light-induced flight bouts intermixed with freezing episodes, partly driven by projections onto GAD1^+^ ventrolateral PAG cells (77). Recently, most TH^+^ rostral ZI cells were found to coexpress VGAT (28, 78), and optogenetic manipulation of the GABAergic population within the rostral ZI supports this role. GABAergic ZI cells which project onto dorsolateral and ventrolateral PAG can modulate bi-directionally both the innate and learned flight response (71). Moreover, photoactivation of GABAergic ZI cells projecting to RE suppresses fear generalization and facilitates extinction learning (28). Despite dopaminergic A13 cells overlapping with a larger population of GABAergic ZI cells, they offer some functional dichotomy where photoactivation of A13 cells enhances only extinction recall (28). Additional support for the role of rostral ZI in goal-directed behaviour and appetitive response comes from Zhang and van den Pol (74) who found increased VGAT^+^ rostral ZI firing activity and excitatory postsynaptic currents following 24h of food deprivation and in the presence of gut hunger signalling hormone, ghrelin (79). Moreover, photoactivation of GABAergic rostral ZI neurons and terminals led to disinhibition of glutamatergic neurons of the paraventricular thalamus and evoked compulsive eating (74). GABAergic rostral ZI cells are also activated during hunting and photoactivation promotes hunting (73). However, there is a slightly greater effect by photoactivation of GABAergic medial ZI cells that incorporate hunting-related sensory signals into hunting motor response. Photoactivation of GABAergic medial ZI cells can also evoke masseter muscle activity and restore impaired hunting from visual and somatosensory deprivations. GABAergic medial ZI projections to PAG appear to be critical in mediating hunting efficiency and appetitive motivational drive.

The A13 region is preserved in PD and could provide an alternative motor response pathway. The present study examined modular reorganization based on anatomical distributions. Future studies should examine functional recruitment and flexibility within the preserved A13 pathways and their therapeutic benefits. DBS of the caudal ZI has been more effective at treating tremors (80–82) and proposed as an alternative therapeutic target for PD. The medial portion of rostral ZI found in rodents where A13 region resides has yet to be explored in the context of PD, and is quite distant from the caudal ZI targets described in human surgeries which are dorsal/dorsomedial to the standard STN target (80). This corresponds to the central (dorsal and ventral) region of ZI in rats (83). Based on surgical and DBS-mediated lesioning of ZI alleviating motor symptoms of PD (80), ZI activity seems to be altered following nigrostriatal degeneration. Périer and colleagues (35) found that ZI cells are hyperactive in the 6-OHDA rat model using electrophysiological recordings and a metabolic marker, cytochrome oxidase. While upregulated A13 pathways indicate compensatory changes and maybe promising potential alternative targets, excessive synchrony or hyperconnectivity between regions may also be pathological (see reviews: (84–86)). Lench et al. (87) found cortical hyperconnectivity to the MLR is correlated to freezing of gait severity. Increased cortico-mesencephalic functional connectivity during ON-state was related to longer single-task and dual-task performance (87). Patients with freezing of gait show higher high-beta oscillations (21-35 Hz) in the subthalamic nucleus at rest in drug-off state than patients without freezing of gait (88). Increased low-beta (13-20 Hz) oscillations in the local field potential (LFP) of the subthalamic nucleus are associated with tremor onset (89), rigidity and bradykinesia (90). In addition, opposing patterns of compensation such as ipsilesional downregulation and contralesional upregulation of A13 efferents to the motor cortex and the striatum may further exacerbate the imbalanced motor outflow from unilateral nigrostriatal degeneration. Activating these A13 efferents would increase motor dysfunction by producing greater striatal and motor cortical activity within the intact, unlesioned side. Thus, it is important to consider the widespread changes in the A13 connectome as a feature of pathophysiology and as a potential contributor to ongoing disease progression and select A13 pathways that would continue to restore motor output in the presence of aberrant connectivity of surviving circuits.

### Implications for motor control

The canonical dopamine (DA) motor pathway is centred on the nigrostriatal circuit (91). DA-containing cells from the substantia nigra pars compacta (SNc) project onto medium spiny GABAergic neurons of the striatum. SNc DA provides an overall excitatory effect on motor output mediated through basal ganglia outputs (globus pallidus internal (GPi)/substantia nigra reticulata (SNr)) to brainstem locomotor regions and cortical motor areas. In PD, striatal dopamine depletion leads to overactive STN and GPi, increased synchronization between pathways affecting motor cortex (see reviews: (84–86)). Although the control of locomotion by DA neurons is thought to be indirect, multiple lines of evidence now challenge this assertion (92–95). Our work shows that A13 projections are affected at cortical and striatal levels following 6-OHDA, presumably targeting motor function.

### Limitations

Here we used AAV tracers that label all processes of the infected cell. However, there are areas such as the striatum where axons pass through without synapsing and require visual inspection of 2D sections for synaptic boutons. Future studies could replace the cytoplasmic YFP AVV with synaptophysin-EGFP-expressing AAV to confidently make this distinction. At present, it remains challenging to target the A13 region based on neurochemistry. The A13 region lacks DAT expression (16, 96), and the TH-cre driver line has an overexpression in the hypothalamic region (95). While most TH^+^ A13 cells colocalize with VGAT+, it remains hard to delineate VGAT+/TH-cells from the region. To facilitate future translational work applying DBS to this region, we targeted the A13 region using a CaMKII-cre driver line. While 6-OHDA models fail to capture the age-dependent chronic degeneration observed in PD, it provides additional advantages in examining robust motor deficits with acute degeneration and identifying compensatory changes compared to the unlesioned side. There are currently few animal models available, which adequately model both the progression and the full extent of SNc cellular degeneration while meeting the face validity of motor deficits . Nevertheless, our data from a human patient with advanced PD points to the preservation of the A13 region, suggesting that the region could participate in compensatory remodelling of the connectome.

### Conclusions

Parkinson’s disease produces changes in motor, sensory, and cognitive function. For these reasons, it is critical to understand the changes to the connectome beyond the nigrostriatal axis. Here we demonstrate a profound restructuring of afferent and efferent inputs to the A13 region, an area involved in coordinating multisensory input with appropriate visceral, arousal, attention, and motor responses. Our work offers predictions in terms of behavioural compensation, where the A13 region may in PD contribute to pathology, but whereby closed-loop manipulation of bilateral A13 could lead to recovery of function. Combined with other work showing reorganization of other regions following degeneration of the nigrostriatal axis, it also points to an underlying explanation for heterogeneity in the progression and severity of PD.

## Materials and Methods

### Animals

All care and experimental procedures were approved by the University of Calgary Health Sciences Animal Care Committee (ACC19-035). Male C57BL/6 mice 7-8 weeks of age were group-housed (≤ 4 per cage) on a 12-h light/dark cycle with ad libitum access to food and water. Mice were randomly assigned into the groups described in the following experiments. See *SI Appendix* for details.

### AAV Vectors

AAV8-CamKII-mCherry (titer 2×10^13^GC/ml, lot 820) was obtained from Viral Vector Facility (Neurophotonics Facility, Laval University, Quebec City, Canada). AAVrg-CAG-GFP (titer ≥ 7×10^12^vg/mL, category number 37825, lot V9234) was purchased from Addgene (Watertown, MA, USA).

### Surgical Procedures

Mice (7-8 weeks of age, range of wt) were anesthetized using isoflurane (1-2%) and stabilized in a stereotaxic apparatus. Stereotaxic injections were performed using a glass capillary (Drummond Scientific, PA, USA; Puller Narishige, diameter 15–20 mm) and a Nanoject II apparatus (Drummond Scientific, PA, USA). Unilateral microinjections of either 0.239 µL vehicle (0.2% ascorbic acid in 0.9% saline; Tocris) or 0.239 µL 6-OHDA (3.6 µg, 15.0 mg/mL; Tocris) and a 50:50 mix of AAVs (total volume 0.110 µL) were made into the MFB and the A13, respectively. Details are provided in *SI Appendix*.

### Behavioral Tests

Mice were habituated in the testing room for at least 0.5 hour for the following tests. The following tests were performed up to 7 days prior to injections and repeated from 21 days post injections. More detailed information is provided in the SI *Appendix*.

#### Treadmill gait test

Each mouse was placed in a polymethyl methacrylate chamber with a clear treadmill (40.5 cm length by 5.2 cm width, Mouse Specifics Inc., Framingham, MA, USA) which allows the animal to walk or run at a fixed velocity. Paw placements were recorded with a video camera (Fujinon HF16HA-1 TV Lens 1 1.4/16 mm, 166 frames/s) mounted underneath the treadmill. We collected up to 180 s of locomotion at 15 cm/s per trial, and three trials were performed per mouse. The videos were analyzed using Visual Gait Lab software, a custom-designed gait analysis software (97). Briefly, Visual Gait Lab uses DeepLabCut (98), a supervised machine learning algorithm for markerless visual tracking, to track paw placements. The resulting tracked position coordinates can be used to compute different gait metrics.

The ability for mice to sustain locomotion was assessed by deviations from mice remaining stationary relative to the camera. As entropy-based measures have been successfully applied to a variety of time-series data types (39, 40), normalized spectral entropy (Hs_n_), which is derived from fast Fourier transform spectral analysis, was used to assess deviations in the stationarity of mice during the longest stretches of locomotion (>1500 frames) in pre- and post-injection time points. Hs_n_ provides a single output measure that gives information about signal fluctuations’ overall complexity; lower values are associated with large deviations in stationarity resulting from discontinuous bouts of locomotion, and higher values being associated with less deviations in stationarity during continuous treadmill running. ΔHs_n_ represents the difference in Hs_n_ post minus pre-injection.

In addition, the best uninterrupted locomotor bout of at least 3 step-cycles (≈150 frames) was used to compare changes in gait metrics following an injection. VGL software imports tracked position coordinates into a custom Python script to compute gait metrics (97). Footfall patterns and interlimb coordinations were analyzed *post-hoc* by importing VGL outputs into custom Matlab scripts (99).

#### Rotarod test

Mice were trained for up to 5 minutes to maintain walking on the rotarod (LE8500, Panlab S.L.U, Barcelona, Spain) at a constant speed of 4 rpm for at least 30 s. There were three testing trials per mouse with a minimum 1-minute break in between trials. Each trial consisted of the rotarod accelerating from 4 to 40 rotations per minute over 3 minutes. Each trial ended when the mouse fell and triggered a time switch upon landing on the equipment base.

#### Open field test

Each mouse was placed in a 70 cm (width) x 70 cm (length) x 50 cm (height) chamber with opaque walls and recorded for 30 minutes using a vertically mounted video camera. Locomotor performance was analyzed *post-hoc* using TopScan video tracking software (CleverSys Inc., Reston, VA, United States) as described previously (95).

### Immunohistochemistry

Mice were anesthetized with isoflurane (2%) and transcardially perfused with PBS, followed by 4% paraformaldehyde (PFA) in PBS. Brains were removed and postfixed overnight in 4% PFA in PBS at 4°C. The next day, a modified iDISCO method (100) was used to clear the samples and perform quadruple immunohistochemistry (anti-GFP, anti-mCherry, anti-TH, and TO-PRO3), as described in detail in *SI Appendix*.

Hypothalamic and midbrain 6 µm sections of formalin-fixed, paraffin-embedded human brain were obtained. TH immunolabeling with horseradish peroxidase (HRP) - 3,3-diaminobenzidine (DAB) detection and hematoxylin counterstain was performed using an Dako Omnis automated machine (Agilent Technologies, Inc., Santa Clara, CA, USA). *SI Appendix* provides more details.

### Imaging

Cleared whole brain samples were imaged using a lightsheet microscope (LaVision Biotech UltraMicroscope, LaVision, Göttingen, Germany) with a 2x objective and 4x optical zoom. *SI Appendix* provides more details. Human immunostained brain sections were imaged on an Olympus VS110 Slide Scanner microscope (Olympus, Center Valley, PA, USA) using a 40x objective.

### A13 connectome analysis

The images were processed with Image J (101). Segmentation for cell counts was achieved manually using ImageJ, while fibres were segmented semi-automatically using Ilastik (102). Images and segmentations were imported into WholeBrain software to be registered with the 2008 Allen reference atlas (103). To minimize the influence of experimental variation on the total labelling of neurons and fibres, the afferent cell counts or efferent fibre areas in each brain region were divided by the total number found in a brain to obtain the proportion of total inputs and outputs. Connectome analyses were performed using R Studio software. For cross-correlation analyses, the data were normalized to a log_10_ value to reduce variability and bring brain regions with high and low proportions of cells and fibres to a similar scale. Hierarchical clustering of complete Euclidean distance matrices by column (brain regions) were determined for each condition based on Spearman’s correlation. The hierarchical cluster dendrograms were trimmed at the optimal tree-cutting threshold (Fig 6A, D) for each given tree to split into specific modules. *SI Appendix* provides more details.

### Statistical analysis

SPSS version 26.0 (IBM) and GraphPad Prism version 9.1.1 (San Diego, California USA) were used for all statistical analyses. Details are provided in *SI Appendix*.

## Acknowledgments

We would like to acknowledge support from Whelan and Kiss Labs and technical support from Hotchkiss Brain Institute Advanced Microscopy Platform Core Facility, Cumming School of Medicine Optogenetics Platform Core Facility and Drs. David Elliot, Jonathan Epp, and Lothar Resch. We acknowledge studentships from Parkinson Alberta (LHK), Parkinson Canada (LHK), Cumming School of Medicine (LHK), Faculty of Graduate Studies (LHK), and the Faculty of Veterinary Medicine (NM, ST). This research was supported by grants to PJW provided by a Canadian Institutes of Health Research Project Grant, Wings for Life, and NSERC (Discovery grant).

