## Supplementary Data for "Substantia nigra degradation results in widespread changes in medial zona incerta afferent and efferent connectomics"

**Affiliations:** [1]Hotchkiss Brain Institute, University of Calgary, Calgary, AB, Canada, T2N4N1, [2]Department of Neuroscience, University of Calgary, Calgary, AB, Canada, T2N 4N1, [3]Department of Comparative Biology and Experimental Medicine, University of Calgary, Calgary, AB, Canada, T2N4N1, [4]Calgary Laboratory Services, Foothills Medical Centre, Calgary, AB, Canada, T2N 2T9, [5]Department of Clinical Neurosciences, University of Calgary, Calgary, AB, Canada, T2N 4N1

*** Corresponding author**
Patrick J. Whelan
HMRB 168,
3330 Hospital Drive NW,
University of Calgary
Calgary, AB
T2N 4N1

**This PDF file includes:**

Supplementary text: 4 pages

Figures S1 to S2

Tables S1

Legends for Movies S1

Legends for Datasets S1

SI References: 5

**Other supplementary materials for this manuscript include the following:**

Movies S1

Dataset S1

**Supplementary Information Text**
**Materials and Methods**

**Animals** Male C57BL/6 mice 7-8 weeks of age (n=3 for saline treatment, n=8 for 6-OHDA injections, but only 6 survived) were used. This is in line with published reports on 6-OHDA mortality (1). All mice were group-housed (≤4 per cage) on a 12-hour light/dark schedule (07:00 lights on - 19:00 lights off) with ad libitum access to food and water. All procedures were approved by the University of Calgary Health Sciences Animal Care Committee (ACC19-035).

**Adenovirus-associated virus (AVV) vectors** AAV8-CamKII-mCherry (titer $2x10^{13}$ GC/ml, lot 820) was obtained from the Viral Vector Facility (Neurophotonics Facility, Laval University, Quebec City, Canada). AAVrg-CAG-GFP (titer ≥ $7\times10^{12}$ vg/mL, category number 37825, lot V9234) was purchased from Addgene (Watertown, MA, USA).

**Surgical procedure for unilateral 6-hydroxydopamine (6-OHDA) lesions of SNc DA neurons.**
We established a well-validated unilateral 6-OHDA mediated Parkinsonian mouse model (2) (Fig 1, *Movie S1*). Thirty minutes prior to stereotaxic microinjections, mice were intraperitoneally injected with desipramine hydrochloride (2.5 mg/ml, Sigma Aldrich) and pargyline hydrochloride (0.5 mg/ml, Sigma Aldrich) (0.9% sterile saline, pH 7.4) at 10 ml/kg to enhance selectivity and efficacy of 6-OHDA induced lesions (2). All surgical procedures were performed using aseptic techniques and mice were anesthetized using isoflurane (1-2 %) delivered by 0.4 L/min of medical grade oxygen (Vitalair 1072, 100 % oxygen).
Mice were stabilized on a stereotaxic apparatus. Small craniotomies were made above the medial forebrain bundle (MFB) and the A13 nucleus within one randomly assigned hemisphere. Stereotaxic microinjections were performed using a glass capillary (Drummond Scientific, PA, USA; Puller Narishige, diameter 15–20 mm) and a Nanoject II apparatus (Drummond Scientific, PA, USA). 0.239 µL of 6-OHDA (3.6 µg, 15.0 mg/mL; Tocris, USA) was microinjected into the MFB (AP -1.2 mm; ML ±1.1 mm from bregma; DV -5.0 mm from the dura). Sham mice received a vehicle solution instead (0.239 µL of 0.2% ascorbic acid in 0.9% saline; Tocris, USA). For tracing purposes, a 50:50 mix of AAV8-CamKII-mCherry (Viral Vector Facility, Neurophotonics Facility, Laval University, Quebec City, Canada, lot 820, titer $2x10^{13}$ GC/ml) and AAVrg-CAG-GFP (Addgene, Watertown, MA. category number 37825, lot V9234, titer ≥ $7\times10^{12}$ vg/mL) was injected ipsilateral to 6-OHDA injections at the A13 nucleus in all mice (AP -1.22 mm; ML ± 4.0 mm from the bregma; DV -4.5 mm from the dura, the total volume of 0.110 µL at a rate of 23 nl/sec). Post-surgery care was the same for both sham and 6-OHDA injected mice. The animals were sacrificed five weeks after surgery.

**Behavioural Tests** Mice were habituated in the testing room for at least 1 hour for the following tests. The following tests were performed up to 7 days prior to injections and repeated from 21 days post injections (see Fig. 2).

***Treadmill gait tests*** Each mouse was placed in a polymethyl methacrylate chamber with a clear treadmill (polyvinylchloride and high-density polyethylene blend, 40.5 cm length by 5.2 cm width, DigiGait™, Mouse Specifics Inc., Framingham, MA) which allows the animal to walk or run at a fixed velocity. Mice were trained to walk consistently on DigiGait™ (Mouse Specifics Inc., Framingham, MA) for at least 3 step cycles at 15 cm/s. DigiGait™ is a clear treadmill (polyvinylchloride and high-density polyethylene blend, 40.5 cm length by 5.2 cm width) in a polymethyl methacrylate chamber that allows the animal to walk or run at a fixed velocity. A camera (Fujinon HF16HA-1 TV Lens 1 1.4/16 mm, 166 frames/s) was mounted underneath the treadmill to record paw placement. To minimize the stress induced by forced movement, each mouse was habituated in the treadmill chamber for 5 minutes at 0 cm/s. Subsequently, the treadmill speed was gradually increased to 15 cm/s at 1 cm/s steps. Each mouse walked for three trials for a maximum of 30 s long at 15 cm/s with a minimum 30 s break between trials. We collected gait measurements 7 days before and 28 days after the stereotaxic injections. Thus, we collected approximately 180 s of data per mouse. The video was then converted to AVI format and imported into Visual Gait Lab software, a custom-designed gait analysis software ((3)).

***Gait Analysis*** Gait analysis was performed using Visual Gait Lab software, and its method has been published previously (3). Briefly, Visual Gait Lab uses DeepLabCut (4), a supervised machine learning algorithm for markerless visual tracking, to track paw placements. The resulting tracked position coordinates were used to compute different gait metrics. For example, the ability of mice to sustain locomotion was evaluated using a matched (pre/post) longest stretch of locomotion (>1500 frames) of the contralesional hindlimb. This corresponded to n=3 saline and n=4 6-OHDA mice. When mice speed is similar to the treadmill speed, they remain stationary relative to the camera. Thus, the ability of mice to sustain locomotor bouts can be reflected by deviations in this stationarity. Normalized spectral entropy ($Hs_n$), which is derived from fast Fourier transform spectral analysis (5), was used to assess deviations in the stationarity of mice during stretches of longer-duration locomotor bouts. $Hs_n$ provides a single output measure that gives information about the overall complexity of signal fluctuations; lower values are associated with large deviations in stationarity resulting from discontinuous bouts of locomotion, and higher values are associated with fewer stationarity deviations during continuous treadmill running. $\Delta Hs_n$ represents the difference in $Hs_n$ post minus pre-injection.

In addition, various gait metrics within the best uninterrupted locomotor bout of at least 3 step cycles (≈150 frames) were compared pre-and post-injections (n=3 saline and n=6 6-OHDA mice). VGL software imports tracked position coordinates into a custom Python script to compute gait metrics (3). Briefly, the following gait metrics from VGL outputs were analyzed across four limbs: average stance duration, average swing duration, average stride duration, ratio between swing to stance durations, average stance width, average stride length, stride length variability, average paw angle, and gait symmetry between forelimbs stride frequencies compared to hindlimbs stride frequencies. In addition to these metrics, footfall patterns and interlimb coordinations were analyzed post-hoc by importing VGL outputs into custom Matlab scripts (6).
Regularity index was computed multiplying the number of normal step sequence patterns by a factor of 4 and dividing by the total number of paw placements and expressing it as a percentage. Six normal step sequence patterns have been previously described and involve cruciate, alternate, and rotary step patterns (7, 8).

$$Regularity\ index\ = (\#\ of\ normal\ step\ sequence\ patterns\ x\ 4/\ total\ number\ of\ paw\ placements)\ x\ 100\%$$

***Rotarod*** Mice were trained for up to 5 minutes to maintain walking on the rotarod (LE8500, Panlab S.L.U, Barcelona, Spain) at a constant speed of 4 rpm for at least 30 s. Those that failed to learn to walk for at least 30 seconds during the 5-minute training periods were returned to their home cage for a minimum of 30 minutes and then retrained until they could achieve a 30-second walk. There were three testing trials per mouse with a minimum 1-minute break in between trials. Each trial consisted of the rotarod accelerating from 4 to 40 rotations per minute over 3 minutes. Each trial ended when the mouse fell and triggered a time switch upon landing on the base of the equipment or when the protocol was completed.

***Open field test*** Each mouse was placed in a 70 cm (width) x 70 cm (length) x 50 cm (height) chamber with opaque walls and recorded for 30 minutes using a vertically mounted video camera. Post hoc analysis was performed using TopScan video tracking software (CleverSys Inc., Reston, VA, United States), which quantifies behaviours such as movement path, distance, speed, displacement, and rearing (9).

**Immunohistochemistry** Mice were anesthetized with isoflurane (2%) and transcardially perfused with PBS, followed by 4% paraformaldehyde (PFA) in PBS. Brains were removed and postfixed overnight in 4% PFA in PBS at 4°C. The next day, a modified iDISCO method (10) was used to clear the samples and perform quadruple immunohistochemistry. The modifications include prolonged incubation and the addition of SDS for optimal labelling. The protocol is provided in Table S1.

Antibodies used were: chicken monoclonal anti-GFP (1:1000, Aves Lab GFP1020, Tigard, OR), rat monoclonal anti-mCherry (1:500, Invitrogen M11217, Thermo Fisher, Waltham, Massachusetts), rabbit polyclonal anti-TH (1:500, Abcam AB112), TO-PRO-3 (1:5000, Invitrogen T3605), Alexa Fluor 488 donkey anti-chicken (1:200, Jackson ImmunoResearch 703-545-155, West Grove, PA), Alexa Fluor Cy3 donkey anti-rat (1:200, Jackson ImmunoResearch 712-165-153), and Alexa Fluor 790 donkey anti-rabbit (1:200, Invitrogen A11374).

Hypothalamic and midbrain sections of the formalin-fixed, paraffin-embedded human brain were obtained in 6 μm thickness. TH immunolabeling with HRP-DAB detection and hematoxylin counterstain was performed using a Dako Omnis automated machine (Agilent Technologies, Inc., Santa Clara, CA, USA). Slices were pre-treated using heat mediated (97 °C) antigen retrieval with sodium citrate buffer (pH 6.0). They were then incubated with rabbit polyclonal anti-TH (1:500, Abcam AB112) and visualized using HRP-DAB coupled immunohistochemistry. The slices were counterstained with hematoxylin. All the reagents (including HRP (GV900), DAB (GV925) and hematoxylin (GC808) were purchased ready to use from Agilent Technologies.

**Imaging** Cleared whole brain samples were imaged using a light sheet microscope (LaVision Biotech UltraMicroscope, LaVision, Göttingen, Germany) with a 2x objective and 4x optical zoom. The brain samples were imaged in an ethyl cinnamate medium to match the refractive indices and illuminated by three sheets of light bilaterally. Each light sheet was 5 μm thick with a numerical aperture of 0.156, and the width was set at 30% to ensure sufficient illumination at the centroid of the sample. Laser power intensities and chromatic aberration corrections used for each laser were as follows: 10% power for 488 nm laser, 5% power for 561 nm laser with 780 nm correction, 40% power for 640 nm laser with 960 nm correction, and 100% power for 785 nm laser with 1,620 nm correction. Each sample was imaged coronally in 8 by 6 squares with 20% overlap (10,202 μm by 5,492 μm in total) and a z-step size of 15 μm.

Immunostained human brain slices were imaged on an Olympus VS110 Slide Scanner microscope (Olympus, Center Valley, PA , USA) using a 40x objective.

**A13 connectome analysis** Images were processed using ImageJ software. Raw images were stitched, and then a z-encoded maximum intensity projection across a 90 μm thick optical section was obtained across each brain. 90 μm sections were chosen because the 2008 Allen reference atlas images are spaced out at around 100 μm. Brains with insufficient quality in labelling were excluded from analysis. YFP+ and TH+ cells were manually counted using the Cell Counter Plug-In (ImageJ). mCherry+ fibres were segmented using Ilastik software and quantified using particle analysis in ImageJ. Images and segmentations were imported into WholeBrain software to be registered with the 2008 Allen reference atlas (n=2 saline and n=3 6-OHDA mice). The TO-PRO-3 channel was used as a reference channel to register each section to a corresponding atlas image. ImageJ quantifications of cell and fibre segmentations were exported in XML formats and registered using WholeBrain software. To minimize the influence of experimental variation on the total labelling of neurons and fibres, the afferent cell counts or efferent fibre areas in each brain region were divided by the total number found in a brain to obtain the proportion of total inputs and outputs. Connectome analyses were performed using custom R scripts (6). For cross-correlation analyses, the data were normalized to a $\log_{10}$ value to reduce variability and bring brain regions with high and low proportions of cells and fibres to a similar scale. Consistency across animals was analyzed using Spearman's correlation (Figure S2). Hierarchical clustering of complete Euclidean distance matrices by column (brain regions) were determined for each condition based on Spearman's correlation. The hierarchical cluster dendrograms were trimmed at the optimal tree-cutting threshold (Fig. 5A,C) for each given tree to split into specific modules.

**Quantification of 6-OHDA mediated TH$^+$ cell loss** The percentage of TH$^+$ cell loss was quantified to confirm 6-OHDA mediated SNc lesions. TH$^+$ cells within ZI, VTA and SNc areas from 90 μm thick optical brain slice images (AP: -0.655 mm to -3.88 mm from bregma) were manually counted by two blinded counters (n=3 saline and n=6 6-OHDA mice; ZI region in 2 of 6 6-OHDA mice were excluded due to presence of abnormal scarring/ healing at the injection site of AAVs). Subsequently, WholeBrain software was used to register and tabulate TH$^+$ cells in the contralesional and ipsilesional brain regions of interest.

Counts obtained from the two counters were averaged per region. The percentage of $TH^+$ cell loss was calculated by dividing the difference in counts between contralesional and ipsilesional sides by the contralesional side count and multiplying by 100 %. Percentage $TH^+$ cell loss = (ipsilesional $TH^+$ cell count – contralesional $TH^+$ cell count)/contralesional $TH^+$ cell count) x 100%.

**Statistical analysis.**
SPSS version 26.0 (IBM) and GraphPad Prism version 9.1.1 (San Diego, California USA) were used for all statistical analyses. Gait metrics obtained using VGL software were analyzed using three-way mixed ANOVA with two within-subjects factors (time and limb types) and one between-subjects factor (treatment type). The time factor consisted of pre-and post-injection measurements. There were two treatment types: saline or 6-OHDA injected. All four limbs of each mouse were referenced to the side of injection: ipsilesional hindlimb, contralesional hindlimb, ipsilesional forelimb, and contralesional forelimb. Gait symmetry and regularity index were analyzed using two-way mixed ANOVA with time as one within-subjects factor and condition as one between-subjects factor. Rotarod performance was analyzed using three-way mixed ANOVA with time and trials as two within-subjects factors and treatment type as one between-subjects factor. Independent t-tests were used to compare percent differences from baseline open field locomotor activity between sham and 6-OHDA mice. Changes in normalized spectral entropy $Hs_n$ post-injections from the baseline values were analyzed between sham and 6-OHDA mice using an independent t-test. Mann-Whitney U test was used when the normality assumption was violated. The means of TH+ cell loss between sham and 6-OHDA mice were compared using two-way mixed ANOVA with brain regions as one within-subjects factor and treatment type as one between-subjects factor. Greenhouse-Geiser corrections were applied when Mauchly's Test of Sphericity failed. Tests of simple main-effects and pairwise multiple comparisons with Bonferroni corrections were performed *post-hoc*.

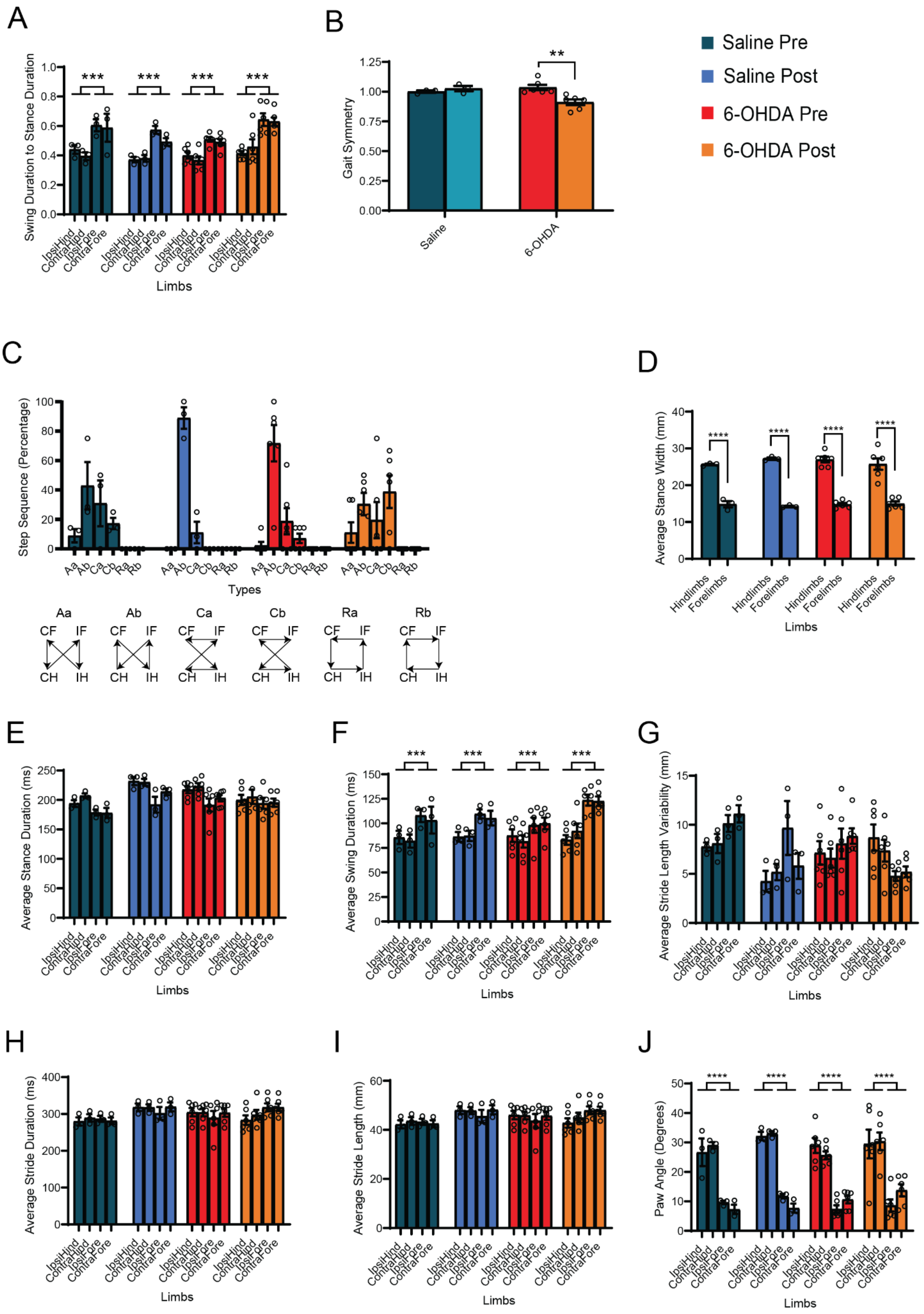
A
B
C
D
E
F
G
H
I
J
Saline Pre
Saline Post
6-OHDA Pre
6-OHDA Post
Swing Duration to Stance Duration
Gait Symmetry
Saline
6-OHDA
Step Sequence (Percentage)
Types
Aa
Ab
Ca
Cb
Ra
Rb
CF
IF
CH
IH
Average Stance Width (mm)
Hindlimbs
Forelimbs
Average Stance Duration (ms)
Average Swing Duration (ms)
Average Stride Length Variability (mm)
Average Stride Duration (ms)
Average Stride Length (mm)
Paw Angle (Degrees)
IpsiHind
ContraHind
IpsiFore
ContraFore
Limbs
***
**
****

**Fig. S1.** Identification of gait abnormalities within one consistent treadmill locomotor bout. There was a significant difference in the swing to stance duration across all four limbs following the 6-OHDA injection (time and condition interaction $p=0.0004$, $F(1,28)=16.53$, A). Gait symmetry between forelimbs to hindlimbs duration was significantly reduced following 6-OHDA injection (time and condition interaction $p=0.010$, $F(1,14)=8.754$, post-hoc $t(14)=4.26$, $p=0.002$, B). In contrast to the sham, 6-OHDA-injected mice demonstrated a significant loss of Ab step sequence dominance (step type, condition and time interaction $p<0.001$, $F(5,42)=11.04$, C). Six normal step sequence patterns in rodents are alternate (Aa and Ab), cruciate (Ca and Cb), and rotary (Ra and Rb) step patterns (7, 8). Other gait features such as lengths and durations of stance, swing or stride did not significantly change over time for either group (D-J, see more details in (3)).

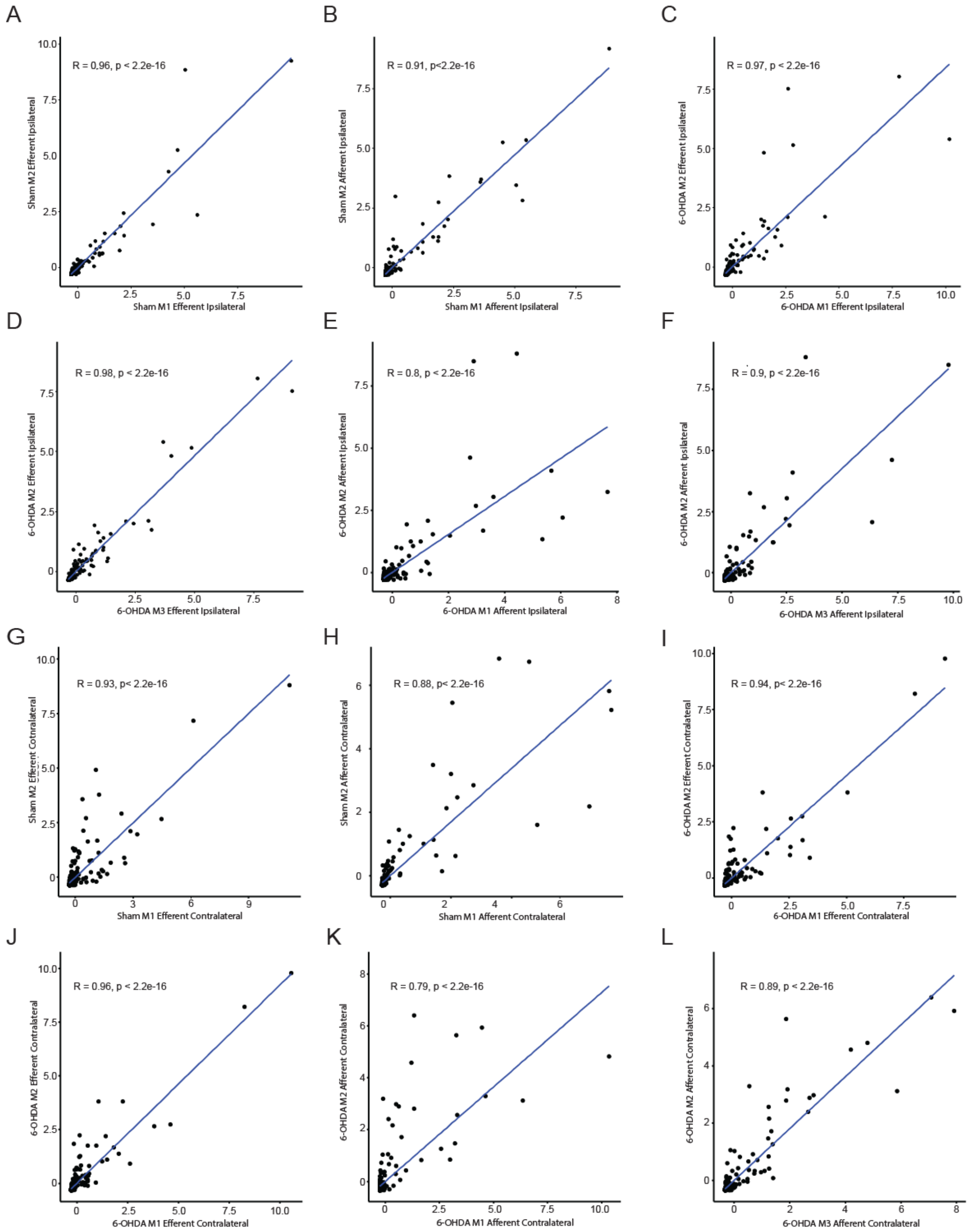


**Fig. S2.** The consistency of afferent and efferent distributions across mice was compared in a pairwise manner. An experimental variation on the total labelling of neurons and fibres was minimized by dividing the afferent cell counts or efferent fibre areas in each brain region by the total number found in a brain to obtain the proportion of total inputs and outputs. Using Spearman's correlation analysis, we found distributions across animals to be consistent among each other with an average correlation of 0.91 (SEM=0.02).

**Table S1**. Modified iDISCO (10) protocol.

| Day # | Instructions |
|---|---|
| 1 | PBS 1x 30 min twice on a shaker at room temperature. |
| 2 | Dehydrate tissue in methanol/H2O series of 20%, 40%, 60%, 80%, 100%, 100% (1 hour each at room temperature) and leave overnight in a 66% DCM/ 33% MeOH solution. |
| 3 | Wash twice in 100% Methanol at room temperature and then chill the sample at 4°c. Bleach the samples in chilled fresh 5% H2O2 in methanol (1 volume 30% H2O2 to 5 volumes MeOH), overnight at 4°C.<br>Before leaving: Bleach in chilled fresh 5%H2O2 in methanol (1 volume 30% H2O2 to 5 volumes MeOH), overnight at 4°c. |
| 4 | Rehydrate in methanol/H2O series of 80%, 60%, 40%, 20%, PBS, PTx.2, PTx,2 (1 hour each at room temperature). Lastly, incubate the samples in Permeabilization Solution at 37°C for 2 days. |
| 6 | Wash 3 times 1-2hr each in 0.5mM SDS/1XPBS, 37°C, then incubate for 3 days. |
| 9 | Incubate with the primary antibody in 0.5mM SDS/1XPBS at 37°C and incubate for 3 days. |
| 12 | Refresh with the primary antibody in PTx.2 at 37°C and incubated for 4 days. |
| 17 | Wash in PTwH for 5 times (2 hours each) and incubate at 37°C overnight. |
| 18 | Incubate with secondary antibody in PTwH/3% Donkey Serum at 37°C for 3 days. |
| 21 | Refresh with secondary antibody in PTwH/3% Donkey Serum at 37°C for 4 days. |
| 25 | Wash in PTwH for 5 times (2 hours each) and incubate at 37°C overnight. |
| 26 | Dehydrate in methanol/H2O series of 20%, 40%, 60%, 80%, 100% (1 hour each at room temperature). Refresh the samples with 100 % methanol and leave overnight at room temperature. |
| 27 | Tissue was incubated for 3 hours in 66% DCM / 33% methanol shook at room temperature. The tissue was then incubated in 100% DCM (Sigma 270997-12X100ML) for 15 minutes twice (with shaking) to wash away methanol. The samples were then incubated with Ethyl Cinnamate for 3 hours at room temperature with shaking. Refresh Ethyl Cinnamate and leave at room temperature for imaging. |

**Movie S1**. A 3D rendition of a unilateral mouse model of Parkinson's Disease. 6-OHDA was injected at the medial forebrain bundle to the lesion nigrostriatal pathway. 3 weeks after, the brain tissue was cleared and immunolabelled using a modified iDISCO protocol. The movie shows immunolabelling against tyrosine hydroxylase antibodies used as a marker for dopaminergic expression. The nigrostriatal pathway of the right hemisphere was lesioned while the left hemisphere remained unlesioned.

**Dataset S1**. Distributions of A13 afferent and efferent expressions across the brain normalized to the total amount of segmentations per brain. SEI = sham efferents ipsilesional side, SEC = sham efferents contralesional side, PEI = PD efferents ipsilesional, PEC = PD efferents contralesional side. SAI = sham afferents ipsilesional side, SAC = sham afferents contralesional side, PAI = PD afferents ipsilesional, PAC = PD afferents contralesional side. Data uploaded to OSF server (6).
